# Analysis of genetic variation in the bovine Mannose Receptor gene (*MRC1)*, its influence on receptor expression, and a potential association with resistance to bovine tuberculosis

**DOI:** 10.64898/2026.06.27.734952

**Authors:** Angela Holder, Jeannine Kolakowski, Eike Usher, Thomas Tzelos, Timothy Connelley, Muhammad Zubair Shabbir, Amanda Gibson, Heather Haris, Bernardo Villarreal-Ramos, Dirk Werling

**Author notes:** Correspondence: Prof. Dirk Werling. Contributed equally to this work (Angela Holder, Jeannine Kolakowski).

## Abstract

Naturally occurring variation in the bovine mannose receptor C-type 1 gene (*MRC1)* may shape macrophage responses to *Mycobacterium (M.) bovis*, a key driver of bovine tuberculosis (bTB). We identified four coding region SNPs in *MRC1* across *Bos taurus* (Holstein Friesian, Brown Swiss) and *Bos indicus* (Boran, Sahiwal) cattle breeds, including a non-synonymous variant, rs380943118 (c.2963G>A; Ser988Asn) in C-type lectin-like domain (CTLD) 6, most prevalent in Sahiwal cattle. Structural modelling suggested that the S988N substitution, which is spatially separated from the monosaccharide binding site of CTLD4, might indirectly affect glycan binding, perhaps through a conformational change in the receptor. Monocyte-derived macrophages upregulated MR expression during differentiation, with heterozygous (G/A) animals showing higher MR expression and increased uptake of GFP-*M. bovis* BCG, although differences were not statistically significant. Anti-CD206 blockade did not inhibit BCG internalization, either indicating that this specific antibody did not bind to a CTLD involved in ligand binding or that MR is not the sole entry receptor. These results highlight naturally occurring *MRC1* polymorphisms that may influence MR structure and macrophage function, providing a foundation for future studies to assess their role in bTB susceptibility.

**Graphical abstract:** 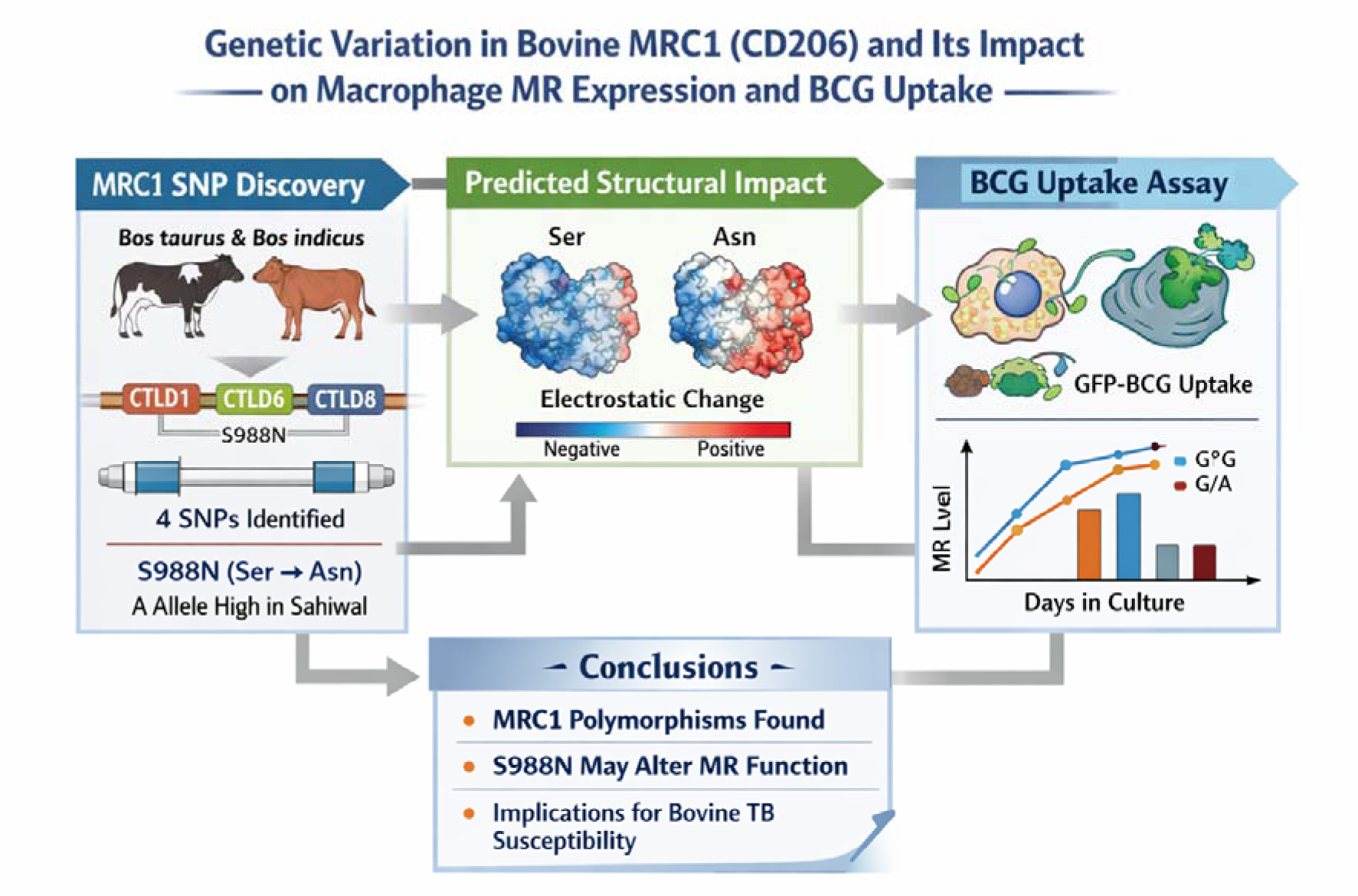

## Introduction

*Mycobacterium tuberculosis* (Mtb) is a member of the *Mycobacterium tuberculosis* complex (MTBC), a group of closely related pathogens responsible for tuberculosis (TB) across a range of mammalian hosts. This complex also includes *M. bovis*, the main causative agent of bovine TB (bTB) and a significant zoonotic pathogen ^1^. Despite ongoing control efforts and enhanced understanding of the molecular mechanisms involved in infection of host target cells, TB remains a major global health burden, with over 10 million human cases reported annually ^2^. Although the majority of TB cases are attributable to infection with Mtb, an estimated 10–15% are caused by *M. bovis* ^3^, underscoring the zoonotic importance of bovine (b)TB and the need for a deeper understanding of host–pathogen interactions that influence susceptibility and disease outcome. *M. bovis* infection costs the global economy an estimated $3 billion annually, with over 50 million infected cattle worldwide ^4^. In the UK alone, managing and eradicating the disease costs approximately £150 million per year in testing, compensation, and research ^5^.

In humans, mycobacterial ligands such as mannosylated lipoarabinomannan (Man-LAM) engage several C-type lectin receptors, including the mannose receptor (MR), facilitating their delivery into early endosomes for antigen processing and presentation via CD1b- and MHC class II–restricted pathways ^6^, and the functional comparison to the bovine MR shows clear similarities ^7^. MR engagement by Man-LAM can also trigger phagocytic uptake of Mtb by macrophages (MØ), although the precise molecular mechanisms governing this process remain incompletely defined ^8^. Notably, MR-mediated internalisation has been associated with inhibition of phagosome–lysosome fusion, thereby enhancing intracellular survival of mycobacteria ^9^.

The Mannose Receptor (MR, CD206), encoded by *MRC1*, is a type I transmembrane C-type lectin receptor expressed predominantly by MØ, dendritic cells (DC), and hepatic and lymphatic endothelial cells ^10^. Its extracellular region is organised into three distinct structural modules that underpin its broad ligand-binding capacity: an N-terminal cysteine-rich domain that recognises sulphated glycans, a fibronectin type II (FnII) domain involved in collagen binding, and eight C-type lectin-like domains (CTLD) that bind glycans terminating in mannose, fructose, or N-acetylglucosamine (GlcNAc) residues ^11–13^. Among these, CTLD4 and CTLD5 serve as the principal carbohydrate recognition domains. While C-type CTLD4 alone is sufficient for monosaccharide binding, high-avidity interactions with complex multivalent glycans depend on the cooperative engagement of CTLD4 to CTLD8 ^14^.

This modular architecture enables the MR to fulfil dual immunological roles. On the one hand, it functions as a scavenger receptor, mediating the clearance of endogenous ligands such as hormones and lysosomal enzymes, thereby contributing to tissue homeostasis ^15^. On the other hand, it acts as a pattern recognition receptor (PRR), detecting glycan structures on a wide range of microorganisms, including yeasts ^16–18^, parasites ^19, 20^, bacteria ^21^, and notably Mtb ^8, 22^. As an endocytic receptor, MR undergoes constitutive internalisation via clathrin-coated pits into early endosomes and is subsequently recycled back to the plasma membrane through a Tyr-Phe di-aromatic motif within its cytosolic tail, thereby avoiding lysosomal degradation ^23^. Consistent with this rapid recycling, only 10–30% of total cellular MR is present at the cell surface under steady-state conditions, with the majority residing intracellularly ^11^. Ligand binding and release are regulated by pH-dependent conformational changes as the receptor traffics between the cell surface and endosomal compartments ^24^.

In line with its established role in pathogen recognition, the MR also serves as an entry pathway for *M. tuberculosis* into human MØ ^25^, and polymorphisms in its encoding gene *MRC1* have been associated with altered susceptibility to human TB ^26^. Although cattle breeds clearly differ in their susceptibility to mycobacterial infection ^27, 28^, the contribution of C-type lectin receptors as early recognition molecules is still poorly understood. Consequently, determining how *MRC1* variation influences MR-mediated recognition and uptake of mycobacteria is essential for identifying host factors that shape resistance or vulnerability to bovine TB.

## Materials and Methods

### Comparative analysis of bovine *MRC1*/MR sequences

The chromosomal location, DNA, and amino acid sequences for bovine *MRC1*/MR (XM_003586772.6) were extracted from the current bovine genome (ARS-UCD2.0) using the NCBI gene database (https://www.ncbi.nlm.nih.gov/). BLAST searches (https://blast.ncbi.nlm.nih.gov/Blast.cgi) of the bovine genome using the nucleotide sequences for human and murine *MRC1* (NM_002438.4 and NM_008625.2) were used to confirm that only one copy of the gene existed. Sequences for sheep, goat, pig, and horse *MRC1*/MR (NM_001197180.1, XM_005687981.3, NM_001255969.1 and XM_005606899.3, respectively) were also obtained from the NCBI gene database for comparison. The percentages of sequence identity were calculated by the pairwise sequence alignment programme EMBOSS Needle (https://www.ebi.ac.uk/Tools/psa/emboss_needle/).

### Samples

Bovine peripheral blood mononuclear cells (PBMCs) were purified by density gradient centrifugation (Lymphoprep, STEMCELL technologies) using whole blood from five Brown Swiss (BS) and five Holstein Friesian (HF) cows reared at the Royal Veterinary College (Hatfield, UK). Experiments were approved under ethical permission PPL7009059, with some restrictions implied. RNA was extracted from boPBMCs using the RNeasy Mini Kit (Qiagen), and cDNA was synthesised using the iScript cDNA Synthesis Kit (Bio-Rad). Complementary DNA (cDNA) samples from five Boran cattle, reared on farms in Africa, and five Sahiwal cattle, reared on farms in Pakistan, were provided by The Roslin Institute (University of Edinburgh, Edinburgh, UK).

### Sequencing and analysis of the bovine *MRC1* gene

Specific primers to amplify the coding region of bovine *MRC1* were designed using nucleotide sequence information (XM_003586772.6) from the NCBI data base and Primer Blast (https://www.ncbi.nlm.nih.gov/tools/primer-blast/) [Suppl. Table 1]. PCR reactions were carried out using Phusion High-Fidelity Master Mix (Thermo Fisher Scientific). PCRs were performed in 50 µl reactions containing 25 µl of 2X Phusion Master Mix, 2 µl of cDNA and 2.5 µl of each primer (10 µM working concentration). Thermocycling conditions consisted of an initial step at 98°C for 30 s, followed by 36 cycles of 98°C for 10 s (denaturation), 65°C or 62°C for 20 s (annealing) and 72°C for 30 s (extension), with a final extension step at 72°C for 10 min (Mastercycler Pro S, Eppendorf). Amplified PCR products were separated by agarose gel electrophoresis, purified using the QIAquick Gel Extraction Kit (Qiagen), and submitted for sequencing (MRC PPU DNA Sequencing Service, Dundee University).

SNPs were identified using the CLC Main Workbench V6.0.2 software (CLCbio, Aarhus, Denmark). Pre-existing dbSNP identifiers were obtained from the European Variation Archive (https://www.ebi.ac.uk/eva/) and Ensembl (https://www.ensembl.org/index.html). Allele and genotype frequencies were calculated for each cattle breed and compared using Fisher’s Exact Test (http://vassarstats.net/index.html).

### Assessment of the structural location of the amino acid affected by SNP rs380943118

The nucleotide sequence for bovine *MRC1* (XM_003586772.6) as well as the amino acid sequence for bovine *MRC1/*MR (XP_003586820.2) were obtained from the NCBI database. 3D protein structures for the protein and the location of SNP rs380943118 (c.2963G>A, S988N) were predicted by AlphaFold 3 Server and analysed using the ChimeraX-1.10.1 software.

### Cell culture

Cells were cultured in tissue culture medium (TCM) which consisted of RPMI-1640 + GlutaMAX-1 (Gibco) + 10% Foetal Bovine Serum (FBS, Sigma-Aldrich) + 1% Penicillin-Streptomycin (PenStrep, Gibco). Monocyte-derived macrophages (MDMØs) were generated from boPBMCs using monocytes (Mo) isolated by either magnetic-activated cell sorting (MACS) or adherence to plastic. For MACS separation, boPBMCs were incubated with anti-CD14 antibody-labelled paramagnetic beads (TÜK4, MicroBeads, Miltenyi Biotec) to isolate CD14^+^ monocytes by positive selection on LS columns in a MidiMACS magnetic separator (Miltenyi Biotech) following manufacturer’s instruction. For adherence to plastic, boPBMCs were incubated in TCM at 37°C and 5% CO_2_ for 24-48 h before non-adherent cells were washed off carefully using pre-warmed 1 x PBS. Adherent monocytes were cultured at approximately 1 x 10^6^ cells mL^-1^ in TCM containing 20 ng mL^-1^ recombinant bovine (rbo) M-CSF (Kingfisher Biotech). On day two post stimulation, remaining non-adherent cells were carefully washed off and spent media was replaced with fresh TCM containing rboM-CSF. After five days in culture, MDMØs were harvested using Accutase (Gibco) according to the manufacturer’s instruction.

### Flow cytometric analysis of MR expression by bovine MDMØ

MDMØ were labelled with a directly conjugated, MR-specific antibody (mouse anti-human CD206-PE, clone 3.29B1.10, Beckman Coulter). Surface staining was analysed on unfixed cells, while total expression was measured using cells that had been fixed and permeabilised (Cytofix/Cytoperm™ kit, BD Biosciences). Cells were blocked prior to antibody staining in 1 x PBS with 1% BSA and 5% mouse serum (Sigma Aldrich) for 20 min at 4°C. Staining was conducted in 1 x PBS with 1% BSA and 1% mouse serum (unfixed cells) or BD Perm / Wash buffer™ (fixed / permeabilised cells) for 40 min at 4°C. Washing was conducted in 1 x PBS with 1% BSA (unfixed cells) or BD Perm / Wash buffer™ (fixed / permeabilized cells). Specificity controls included unstained cells and cells labelled with a corresponding isotype control (mouse IgG1 negative control RPE, MCA928PE, Bio-Rad). Viability was assessed using trypan blue exclusion, which showed less than 10% of dead MDMØ. Flow cytometric analysis was carried out using either a FACSCalibur (BD Biosciences) equipped with two lasers (488nm, 635nm) and standard filters (EP: 488/10, 530/30, 585/42 and 650LP), or a BD LSRFortessa X-20 (BD Biosciences) equipped with four lasers (355nm, 405nm, 488nm, 640nm) and standard filters (EP: 379/28, 450/50, 488/10, 525/50, 530/30, 575/25, 610/20, 670/30, 710/50, 730/45, 780/60). Acquisition settings were determined using the specificity controls, counting 10,000 events that were recorded with the CellQuest Pro software (FACSCalibur, BD Biosciences) or the BD FACSDiva V9 software (LSRFortessa X-20, BD Biosciences). Data were analysed using the FlowJo V9 or V10 software (BD Biosciences). An example of the gating strategy for the analysis of MR expression by bovine MDMØ is shown in [Suppl. Figure 1]. Data generated included percent positive staining and median fluorescence intensity (MFI) ratio (sample/control). Group means were compared by an unpaired t-test with Welch’s correction using the GraphPad Prism V10 software. Comparison of MR surface expression on bovine MDMØ after five days in culture showed no difference between MACS-sorted cells and MDMØ generated via adherence to plastic [Suppl. Figure 2]. The presence of MDMØ was confirmed by flow cytometric analysis of macrophage-expressed surface markers [Suppl. Figure 3] using antibodies listed in [Suppl. Table 2]. Cells were stained and analysed as described earlier in this paragraph.

### Preparation of *M. bovis* Bacillus Calmette-Guerin

The Pasteur strain of *M. bovis* Bacillus Calmette-Guerin (BCG) expressing green fluorescent protein (GFP) was cultured for 14 days at 37°C in Middlebrook 7H9 medium (Difco, BD) containing 10% Middlebrook ADC supplement (Difco, BD), 0.05% Tween80 (Sigma-Aldrich), 2% glycerol and 20 µg mL^-1^ kanamycin. Aliquots were stored at -20°C until further use. Upon defrosting, the BCG was centrifuged at 400 x *g* for 10 min and the pellet resuspended in 1 x PBS by vortexing. Colony forming units (CFU) were estimated from OD_600_ measurements (OD_600_ 1.0 = 3.13 x 10^7^ CFU mL^-1^) and used for subsequent calculation of the MOI ^29^.

### Phagocytosis assay

Bovine MDMØ were seeded in 96-well U bottom plates (3 x 10^5^ cells per well in 200 µl TCM). Cells were incubated with either TCM alone, 1 µg of a mouse anti-human CD206 antibody (clone 3.29B1.10, Beckman Coulter), or 1 µg of a corresponding isotype control (mouse IgG1 negative control, Bio-Rad) at 37°C and 5% CO_2_ for 25 min. Cells were then co-cultured with live GFP-BCG (MOI of 5) in 1 x PBS at 37°C and 5% CO_2_ for 90 min. Cells incubated at 4°C served as control since phagocytosis was reduced under these conditions. Two washes with ice-cold 1 x PBS were followed by killing of surface-bound bacteria via incubation with 50 µg mL^-1^ gentamicin (Sigma-Aldrich) for 30 min. The cells were washed again with ice-cold 1 x PBS, fixed with 4% paraformaldehyde (Cytofix, BD) for 30 min at 4°C, and analysed by flow cytometry using a BD LSRFortessa X-20 (BD Biosciences). Acquisition settings were determined using the specificity controls, counting 10,000 cells that were recorded with BD FACSDiva V9 software (BD Biosciences). Data were analysed using FlowJo V10 software (BD Biosciences). An example of the gating strategy for the analysis of GFP-BCP uptake by bovine MDMØ is shown in [Suppl. Figure 4]. Sample MFI values were normalised by division with control MFI values obtained from MDMØ incubated with GFP-BCG at 4°C. The different treatments were compared by one-way analysis of variance with Geisser-Greenhouse correction adjusted for multiple comparisons with the Tukey test. GFP-BCG uptake by MDMØ with different genotypes for SNP rs380943118 (G/G or G/A) was compared by two-way analysis of variance with Geisser-Greenhouse correction adjusted for multiple comparisons with the Šídák test GraphPad Prism V10 software.

### Analysis of MR antibody binding specificity

The binding specificity of the MR antibody used in this study (mouse anti-human CD206-PE, clone 3.29B1.10, Beckman Coulter) was assessed by ELISA. Recombinant biotinylated fragments (kindly provided by Prof Kurt Drickamer, Imperial College London) for the bovine MR extracellular domain (ECD), CTLD4/5, and CTLD4 alone were tested. Biotinylated human MR (AcroBiosystems) was used as a positive control, while biotinylated bovine Mincle CTLD (kindly provided by Prof Kurt Drickamer, Imperial College London) was used as a negative control. Streptavidin-coated wells (Pierce) were incubated overnight at 4°C with biotinylated protein fragments (1 µg mL^-1^, 100 µl) in assay buffer containing PBS pH7.4 (Gibco), 1 mM CaCl_2_ (Sigma-Aldrich), 0.1% BSA (Sigma-Aldrich). Each protein fragment was assayed in duplicate. All subsequent steps were conducted in assay buffer supplemented with 0.1% Tween20. After being washed three times (200 µl), wells were incubated with the MR antibody (1:50, 100 µl) for 1 h at room temperature, before further washing and incubation with a secondary goat anti-mouse IgG (H+L): HRP antibody (1:2000, 100 µl, Bio-Rad, Cat No. 1706516). The wells were developed with TMB substrate (100 µl, ThermoScientific) and the assay was stopped with 2 N Sulphuric Acid (50 µl, R&D Systems). Absorbance values were measured at 450 nm on an ELISA plate reader (Multiscan FC, ThermoScientific). Background absorbance, measured in duplicate wells that were incubated with the protein fragment and the secondary antibody, was subtracted.

## Results

### Comparative analysis of *MRC1*/MR sequences

Analysis of the current bovine genome (ARS-UCD2.0) identified the sequence for bovine *MRC1* (MR, CD206) as a single copy gene located on chromosome 13 in a region devoid of other C-type lectin receptor genes. The same has been observed for both human and mouse *MRC1* located on the corresponding chromosomes 10 and 2, respectively. Comparison of amino acid sequences of bovine *MRC1*/MR with those of human, mouse, and other livestock species (sheep, goat, pig, and horse) showed a high degree of sequence identity [Table 1]. As expected, comparison of bovine *MRC1*/MR with other ruminant species such as sheep and goat yielded the highest percentage of sequence identity (> 97%).

### Analysis of polymorphisms in the bovine *MRC1* gene

Sequencing of the *MRC1* coding region in cDNA samples from two *Bos taurus* cattle breeds (Brown Swiss n=5, Holstein Friesian n=5) identified four SNPs [Table 2]. All of these SNPs had been identified previously and their details catalogued in the European Variation Archive. (https://www.ebi.ac.uk/eva/). Three SNPs were synonymous (rs110460494, rs208334503, rs109104237) while the fourth (rs380943118) resulted in a serine to asparagine amino acid substitution (S988N). None of the SNPs were found to occur at significantly different frequencies between the BS and HF cattle [Table 3 and 4]. Therefore, the decision was made to investigate the non-synonymous SNP (rs380943118) in two *Bos indicus* cattle breeds (Boran n=5, Sahiwal n=5) [Table 4]. The alternative A allele was found to occur at a significantly higher frequency in Sahiwal cattle compared to BS and HF (p = 0.02). No significant difference was found between HF, BS, and Boran cattle, or between Sahiwal and Boran cattle. Individual cattle homozygous for the alternative A allele were found only in the *Bos indicus* cattle.

### Assessment of the structural location of the amino acid affected by SNP rs380943118

The structural location of the amino acid affected by SNP rs380943118 (S988N) was assessed in 3D protein models of bovine MR CTLDs 4-8 (AA628-1373). The models were generated with AlphaFold 3 Server and analysed using the ChimeraX-1.10.1 software. Published structures of CTLD4 of human MR ^12^ were used to identify the amino acid residues forming the principal Ca^2+^, as well as primary and extended monosaccharide binding sites in the bovine CTDL4 protein model [Figure 1B + C, green and teal residues, respectively]. In comparison, the amino acid residue affected by SNP rs380943118 is in position 988 in CTLD6 [Figure 1B + C, lime green or magenta residue], which is spatially separated from the monosaccharide binding site of CTLD4.

**Figure 1.**
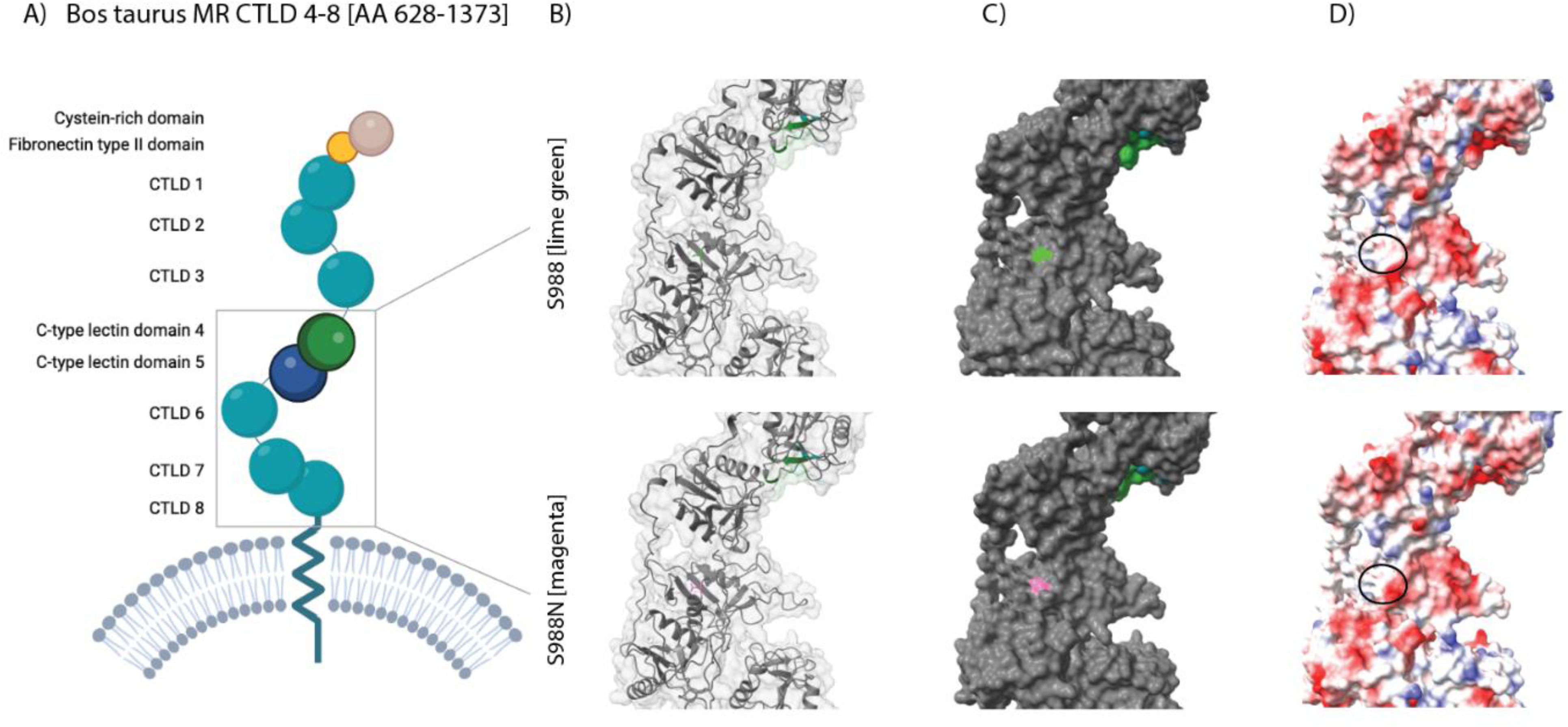
Assessment of the structural location of the amino acid affected by SNP rs380943118. CTLDs 4-8 (amino acid residues 628-1373) of bovine MR were modelled using AlphaFold 3 Server and analysed with the ChimeraX-1.9 software. Amino acid residues forming the principal Ca^2+^ and primary monosaccharide binding site (E743, N745, Y747, Q748, E751, N765, D766, I767) as well as the extended monosaccharide binding site (L712, E755, K757) of CTLD4 in human MR were localised in the bovine models and highlighted in green and teal, respectively. The amino acid residue in CTLD6 of bovine MR that is affected by SNP rs380943118 is shown in lime green (S988, top row) and magenta (S988N, bottom row). The change from ser to asn results in a different electrostatic surface potential in that area (black circle, negatively charged areas in red, positively charged areas in blue).

### MR expression in bovine MDMØ

Expression of MR was examined during the maturation of bovine Mo to MDMØ. Mo were purified from boPBMCs and differentiated into MDMØ by addition of recombinant bovine M-CSF over the course of five days. Surface expression of MR, measured using flow cytometry, was detected on Mo from all the cattle examined and increased during the maturation to MDMØ [Figure 2]. Although expression varied between individual animals during the time course (e.g. 4.35% to 77.62% on day 2), MDMØ cultured with M-CSF for five days were shown to have the highest percentage of MR expression and additional characteristics demonstrating a MØ phenotype. MØ exhibited classical morphological features indicating maturation upon examination by microscopy: compact nucleus, spread out granular cytoplasm compared to Mo. In addition, the cells concurrently showed increase expression of the MØ markers CD16, CD163, and MHC II while continuing to express CD14 on their surface [Suppl. Figure 3].

**Figure 2.**
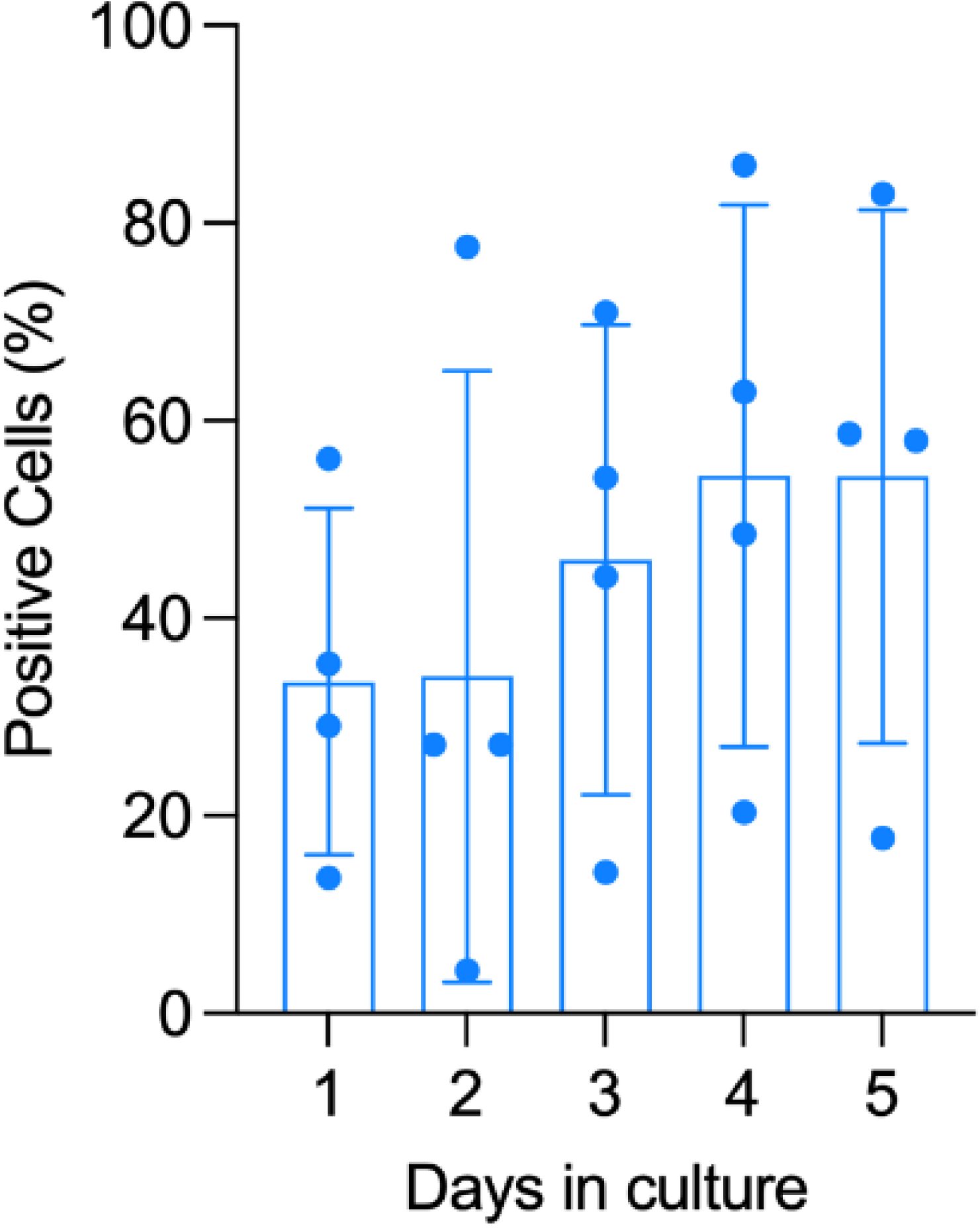
Flow cytometric analysis of MR surface expression on maturing bovine Mo. Bovine monocytes (Mo) isolated from PBMCs of four cattle (1 HF, 3 BS) were differentiated into MØ by addition of recombinant bovine M-CSF for five days. The percentage of maturing Mo expressing MR on their surface was analysed by flow cytometry on each of these five days. Values for individual cattle are shown as filled circles. Data are presented as Means± SD.

Day 5 MDMØ from different *Bos taurus* breeds (BS and HF) were then examined by flow cytometry for both, surface and total expression of the MR [Figure 3]. Evaluation of total expression of MR was performed on fixed and permeabilised cells, taking into account the internalisation and recycling of the receptor known to occur as part of endocytic function. A comparison of the surface and total expression confirms the presence of internalised receptor and suggests approx. 70 -75% of the receptor is on the surface [Figure 3A+F]. Bovine MDMØ from HF and BS cattle showed no breed-specific differences in either surface or total MR expression when analysing percent positive cells (surface P=0.826, total P=0.599) [Figure 3B+C] or MFI ratio (surface P=0.834, total P=0.237) [Figure 3G+H].

**Figure 3.**
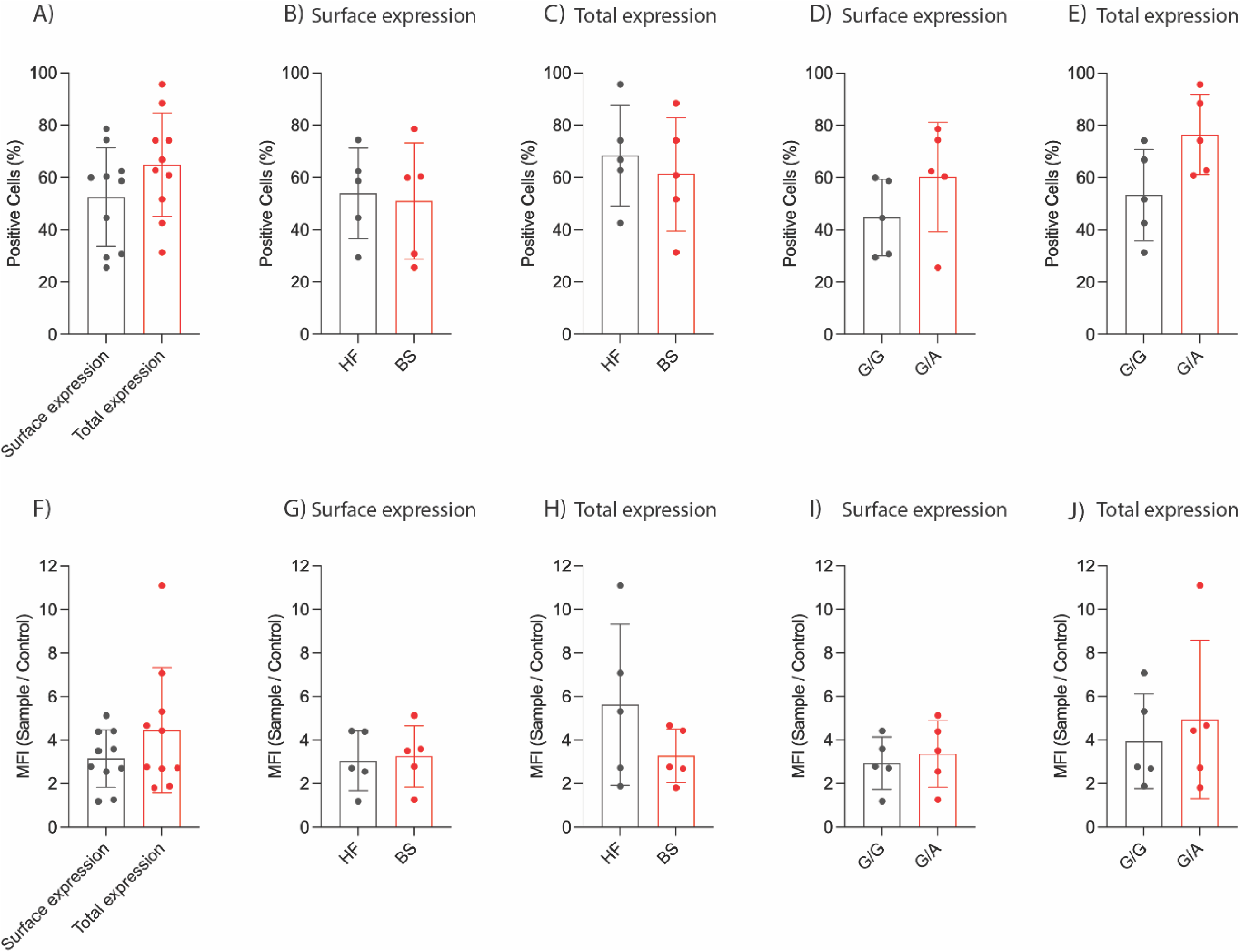
Flow cytometric analysis of MR expression by bovine MDMØ. Bovine monocytes were isolated from PBMCs and differentiated into MØ by addition of recombinant bovine M-CSF for five days. A + F) Unfixed as well as fixed and permeabilised MDMØ were analysed by flow cytometry for surface and total expression of MR, respectively. No statistically significant differences in the percentage of both surface and total MR expression were observed with regards to B + C) cattle breed and D + E) genotype. Equally, no statistically significant differences were observed in the amount of surface and total MR expression with regards to G + H) cattle breed and I + J) genotype. Values for individual cattle are shown as filled circles. The mean is represented by bars and the SD by error bars. MFI ratios were calculated by dividing each sample MFI by the corresponding control MFI. Group means were compared using unpaired t tests with Welch’s correction.

Analysis of the flow cytometry data by SNP rs380943118 genotype showed that heterozygous G/A cattle had a higher average percentage of MR-positive MDMØ than homozygous G/G cattle for both surface and total staining [Figure 3D+E]. However, these differences were not statistically significant (surface P=0.214, total P=0.058), likely due to the small sample size. No differences between G/A and G/G cattle were observed for the MFI ratio (surface P=0.637, total P=0.615) [Figure 3I+J].

### Analysis of BCG-GFP uptake by bovine MDMØ

The MR has been described to be involved in the uptake or phagocytosis of *Mycobacterium spp.* We therefore used *M. bovis* BCG stably expressing green fluorescent protein (GFP-BCG) to assess potential genotype-specific differences caused by SNP rs380943118 in the bovine MR and its role in the uptake activity of bovine MDMØ. Cells from six cattle, three homozygous G/G and three heterozygous G/A, were incubated with GFP-BCG [Figure 4]. Baseline uptake, which was used to normalise the data, was assessed via incubation of cells with GFP-BCG at 4°C. Flow cytometric analysis showed a higher MFI of uptake for MDMØ from G/A heterozygous cattle compared to cells from homozygous G/G cattle [Figure 4B]. However, the difference in uptake between genotypes was not statistically significant under any of the conditions tested (GFP-BCG only P=0.755, isotype control P=0.825, and anti-CD206 Ab P=0.539). Furthermore, the chosen antibody did not seem to affect the level of GFP-BCG uptake by MDMØ (adjusted p value of 0.544) [Figure 4A+B].

**Figure 4.**
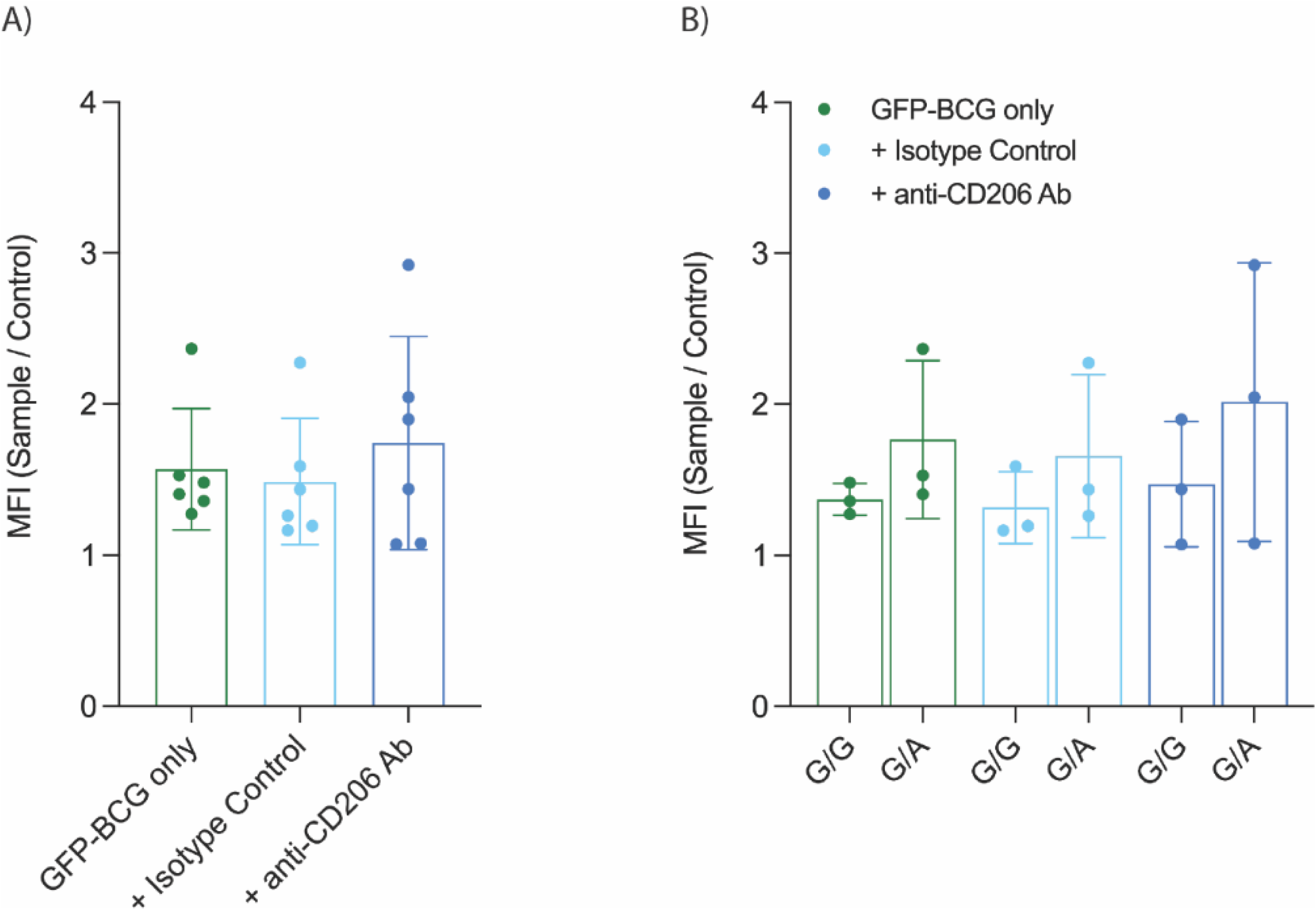
Analysis of BCG-GFP uptake by bovine MDMØ. Bovine MDMØ were incubated with GFP-expressing BCG and analysed by flow cytometry to determine the uptake activity of bovine MDMØ with (G/A) and without (G/G) SNP rs380943118. A) Preincubation of MDMØ with 1 µg of mouse anti-CD206 antibody or a corresponding isotype control had no statistically significant effect on GFP-BCG uptake. B) There was no significant difference in GFP-BCG uptake by MDMØ generated from either G/A or G/G SNP rs380943118. Sample MFI values were normalised via division by control MFI values obtained from MDMØ incubated with GFP-BCG at 4°C. The mean for each condition is represented by bars and the SD by error bars. Different treatments were compared by one-way analysis of variance with Geisser-Greenhouse correction adjusted for multiple comparisons with the Tukey test. GFP-BCG uptake by MDMØ with (G/A) or without (G/G) SNP rs380943118 was compared by two-way analysis of variance with Geisser-Greenhouse correction adjusted for multiple comparisons with the Šídák test.

### Analysis of MR antibody binding specificity

An ELISA using biotinylated MR fragments (ECD, CTLD4, and CTLD4&5) showed that the mouse anti-human CD206 antibody used in this study cross-reacts with bovine MR [Figure 5]. However, the specific epitope recognised by the antibody was found not to be in the two main carbohydrate recognition domains (CTLD4 and 5), but somewhere else in the ECD.

**Figure 5.**
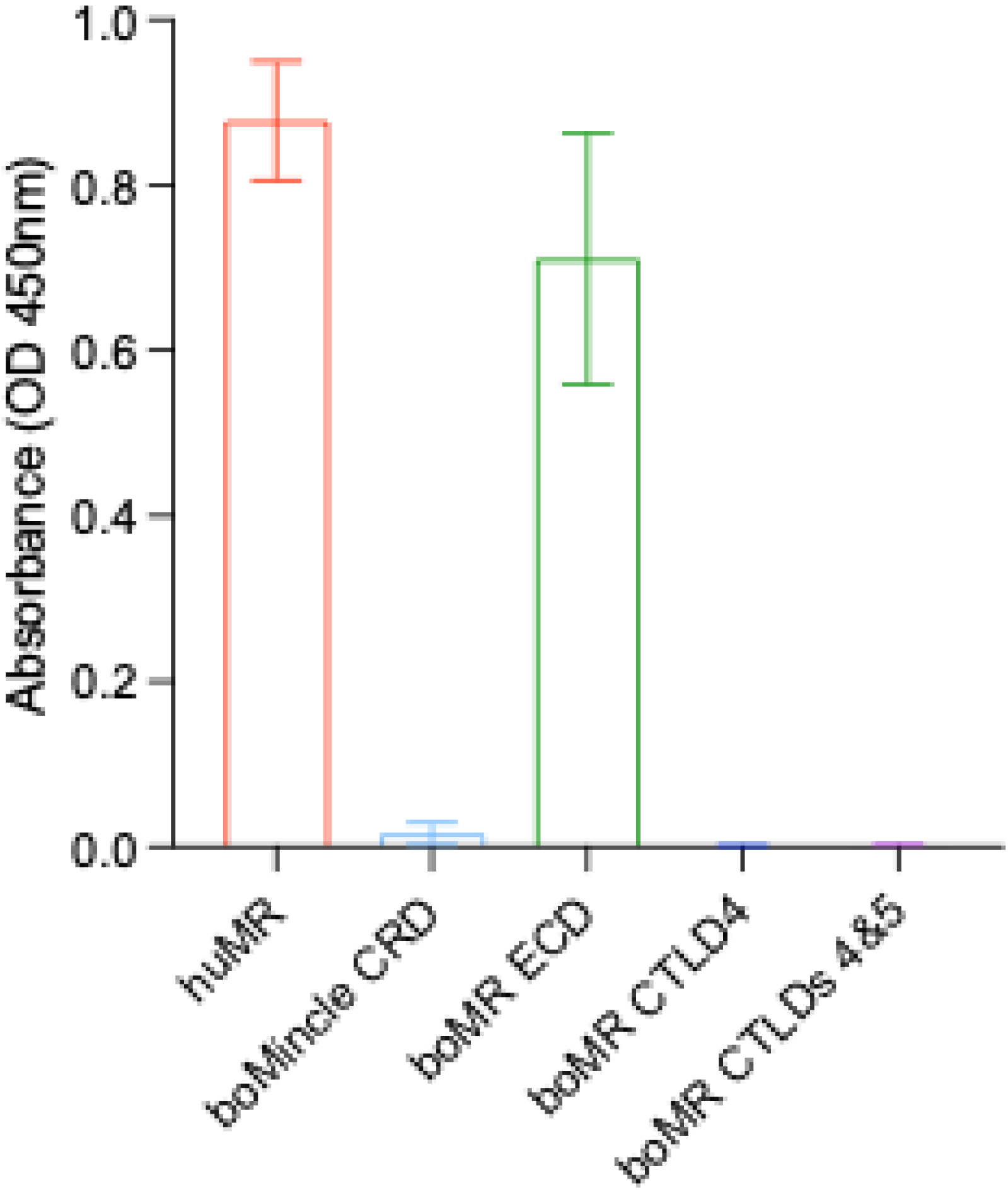
Analysis of MR antibody binding specificity. The mouse anti-human CD206 antibody (clone 3.29B1.10) was tested for cross-reactivity against recombinant biotinylated MR fragments by ELISA. The mean for each condition is represented by bars and the SD by error bars. Human MR and bovine Mincle were included as positive and negative controls respectively. The antibody was found to cross-react with the extracellular domain of bovine MR.

## Discussion

The mannose receptor (MR, CD206) is a C-type lectin receptor (CTLR) within the mannose receptor family that includes further CTLRs such as DEC205, Endo180, and the phospholipase A2 receptor (PLA2R; ^11^). The bovine *MRC1* gene maps to chromosome 13 in a region lacking other CTLR genes, mirroring the dispersed chromosomal organization of members of this family in humans ^11^. Sequence comparisons indicate that bovine MR retains the canonical modular architecture—N-terminal cysteine-rich domain, fibronectin type II domain, eight C-type lectin-like domains (CTLDs), a transmembrane segment, and a short cytoplasmic tail—and shares >80% amino-acid identity with human and murine MR, consistent with conserved ligand recognition.

Polymorphisms in human *MRC1* have been associated with susceptibility to *Mycobacterium* infections, including tuberculosis and leprosy ^26, 30^. We therefore investigated bovine *MRC1* variation and its functional impact. Sequencing of the MR of BS and HF cattle identified four coding SNPs, of which only rs380943118 was non-synonymous. Despite reports of enhanced pathogen resistance in BS relative to HF ^31, 32^, no allele or genotype frequency differences were detected between these breeds, noting the limited sample size. The alternative allele of rs380943118 was most frequent in the Sahiwal of *Bos indicus* that is considered relatively resistant to bovine tuberculosis and least frequent in HF, which is considered to be a susceptible species ^27, 28^. Homozygous carriers of the alternative A allele were observed only within the *Bos indicus* breeds (Boran, Sahiwal).

The MR recognizes host glycans and microbial ligands, including mannose-capped lipoarabinomannan. CTLD4 and CTLD5 account for most carbohydrate binding while only C-type CTLD4 can bind monosaccharides on its own ^14^. While CTLD5–CTLD6 do not bind carbohydrates, CTLD5–7 and CTLD5–8 exhibit carbohydrate binding, but substantially lower compared to CTLD4–5 (Taylor, Bezouska et al., 1992). The rs380943118 variant results in a Ser→Asn substitution at residue 988 in CTLD6. Although CTLD6–8 lack intrinsic monosaccharide-binding activity, they are necessary to support high-avidity interactions with multivalent glycans ^14^. Structural modelling of bovine MR CTLD4–8 demonstrated that the S988N substitution is spatially separated from the main carbohydrate binding domain of CTLD4 and 5 and therefore unlikely to directly affect ligand binding.

However, structural analyses suggest that the MR can adopt at least two distinct domain architectures, and that a transition from a bent, compact arrangement of its extracellular domains to a more extended, linear conformation may function as a regulatory mechanism governing ligand binding (Napper, Dyson et al. 2001; Llorca 2008; Hu, Shi et al. 2018).

Such a change in conformation might alter the proximity and orientation of CTLD6 in relation to CTLD5 and suggest a possible indirect mechanism for SNP rs380943118 to affect ligand binding.

N-glycosylation has also previously been shown to modulate MR glycan binding ^33^, but the lack of an N-X-S/T motif at residues 988–990, argues against the creation of a novel glycosylation site at the position of the SNP.

MR functions predominantly as an endocytic receptor that undergoes constitutive internalization and recycling ^10^. Surface expression is cytokine-regulated: IL-4, IL-10, and IL-13 upregulate MR, whereas IFN-γ induces downregulation ^34–38^. In our MDMØ cultures, ∼70% of total MR was detected at the cell surface, consistent with activation induced during M-CSF-driven differentiation; M-CSF increases IL-10 production in human MDMØ ^39^. MR expression did not differ between BS and HF MDMØ, aligning with the lack of breed-specific coding differences. Heterozygous (G/A) animals showed a higher percentage of MR-expressing MDMØ than homozygous (G/G) animals, although this was not statistically significant.

Bovine MDMØ from heterozygous (G/A) animals internalized more GFP-expressing BCG than MDMØ from homozygous (G/G) animals. This difference was not statistically significant but is in alignment with MR expression. Similar associations between MR levels and phagocytic capacity have been reported in human alveolar macrophages ^40^. However, anti-CD206 (clone 3.29B1.10) did not inhibit BCG uptake. While cross-reactivity of the antibody with bovine MR was confirmed by ELISA, the recognised epitope appeared to be outside of CTLD4 and 5. Prior work indicates stronger functional blockade by the antibody in human than in bovine dendritic cells ^7, 41^, consistent with a conformational epitope and species-specific differences limiting blockade efficacy.

Alternative receptors likely contribute to BCG uptake. A bovine lectin array showed BCG binding to MR, but stronger interactions with DC-SIGN and the ruminant collectins CL-43 and CL-46 ^42^. DC-SIGN is a major receptor for *M. tuberculosis* on dendritic cells ^43–45^, is highly expressed on alveolar macrophages isolated from TB-infected patients ^46^, and can be induced by IL-4 ^47^, and similarly, has been shown to bind *M.bovis*-BCG-GFP in the bovine system ^48^. Given that our MDMØ were not IL-4–polarized, complement receptor 3 (CR3) a constitutively expressed receptor on monocytes and macrophages ^49^ and a well-established mediator of *M. tuberculosis* phagocytosis ^50^ represents a plausible parallel pathway.

In summary, bovine *MRC1* is structurally conserved, and rs380943118 shows breed-associated distribution suggestive of functional relevance. Trends toward higher MR expression and BCG uptake in heterozygous animals were modest and not statistically significant. These data support a model in which MR contributes to, but does not solely determine, mycobacterial uptake by bovine macrophages. Larger cohorts and mechanistic perturbations (e.g., receptor blockade, knockdown/knockout) will be required to define the contribution of *MRC1* variation to bovine tuberculosis susceptibility.

## Acknowledgements

J.F.K was funded through a Bloomsbury PhD studentship to D.W.. E.U. was funded through a summer studentship to A.H.. AJG, and BVR received combined support from the Sêr Cymru II programme (grant number AU185 awarded to Prof. R. G. Hewinson) and Aberystwyth University. AJG is currently supported by UKRI Future Leaders Fellowship (grant number 38860). We thank Prof Kurt Drickamer and Prof Maureen Taylor, both Imperial College for providing recombinant bovine MR fragments. This work was supported by Biotechnology and Biological Sciences Research Council Grants BB/P005659/1 (to K. D. and M. E. T.), BB/P008461/1 (to D. W.). Furthermore, part of this work was supported by a grant from the Gates Foundation through GALVmed to DW (grant number 1583655).

