## Supplementary material for "Analysis of genetic variation in the bovine Mannose Receptor gene (*MRC1)*, its influence on receptor expression, and a potential association with resistance to bovine tuberculosis": Suppl FIle

**Supplementary Figures**

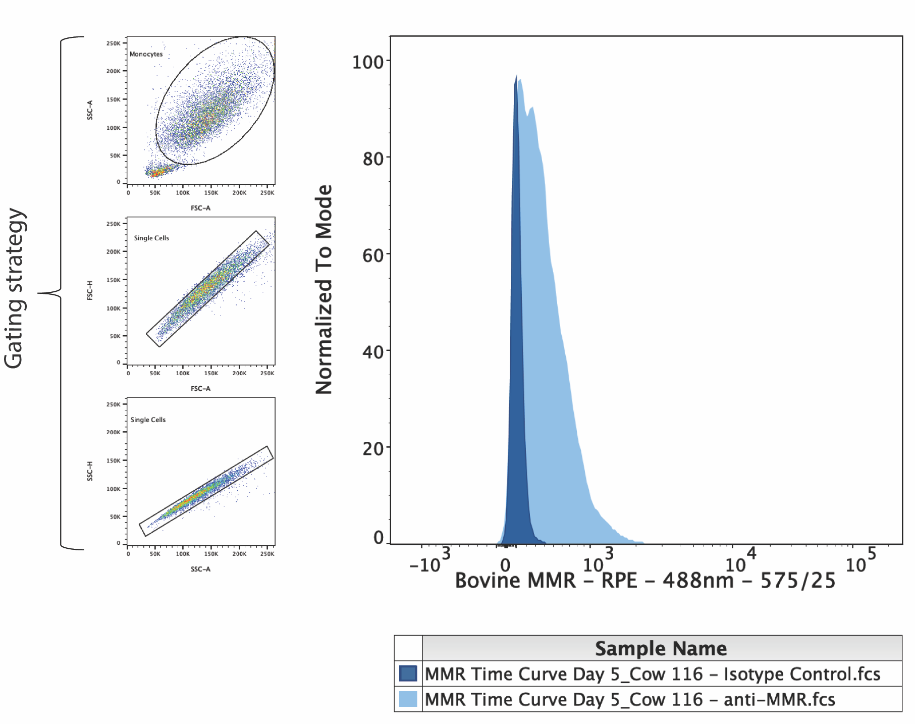

*Supplementary Figure 1 Gating strategy for the analysis of MR surface expression by bovine MDMØ*

*Bovine MDMØ were stained with a directly conjugated anti-CD206 antibody or the corresponding isotype control and analysed by flow cytometry. Detection of all events (FSC vs. SSC) was followed by two doublet discriminations (gating strategy) and assessment of CD206 surface expression on stained cells (light blue histogram) compared to the respective control (dark blue histogram).*

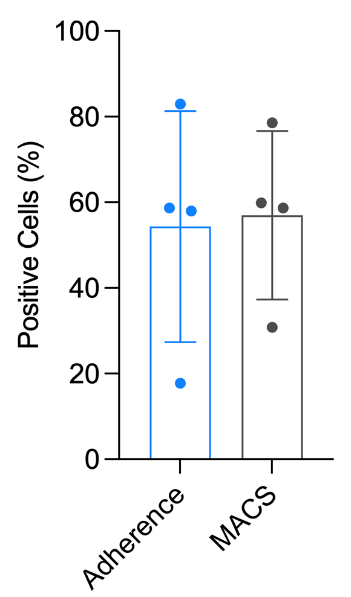

*Supplementary Figure 2 Comparison of MR surface expression by MDMØ generated via adherence to plastic and MACS after five days in culture*

*Bovine Mo* *were isolated from PBMCs via either adherence to cell culture plastic (Adherence) or magnetic activated cell sorting (MACS), and differentiated into MDMØ by addition of recombinant, bovine M-CSF. MR surface expression (MR Positive Cells) was compared via flow cytometric analysis after five days in culture and showed no statistically significant difference between the two methods (unpaired t test with Welch’s correction).*

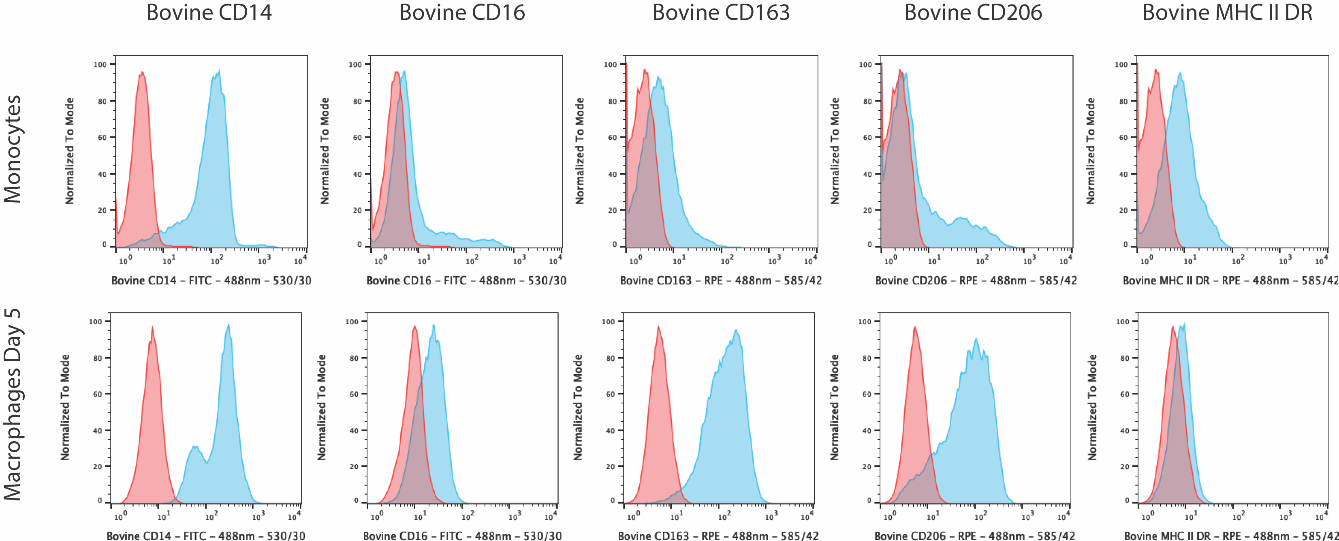

*Supplementary Figure 3 Flow cytometric assessment of bovine Mo differentiation into MDMØ*

*Bovine Mo were purified from PBMCs and differentiated into MDMØ by addition of recombinant, bovine M-CSF. The expected maturation of stimulated Mo was assessed by flow cytometric detection of Mo- and MØ-specific surface markers. The image shows surface staining of Mo and MDMØ after five days in culture (blue histograms) for five of these surface markers compared to the respective isotype controls (red histograms).*

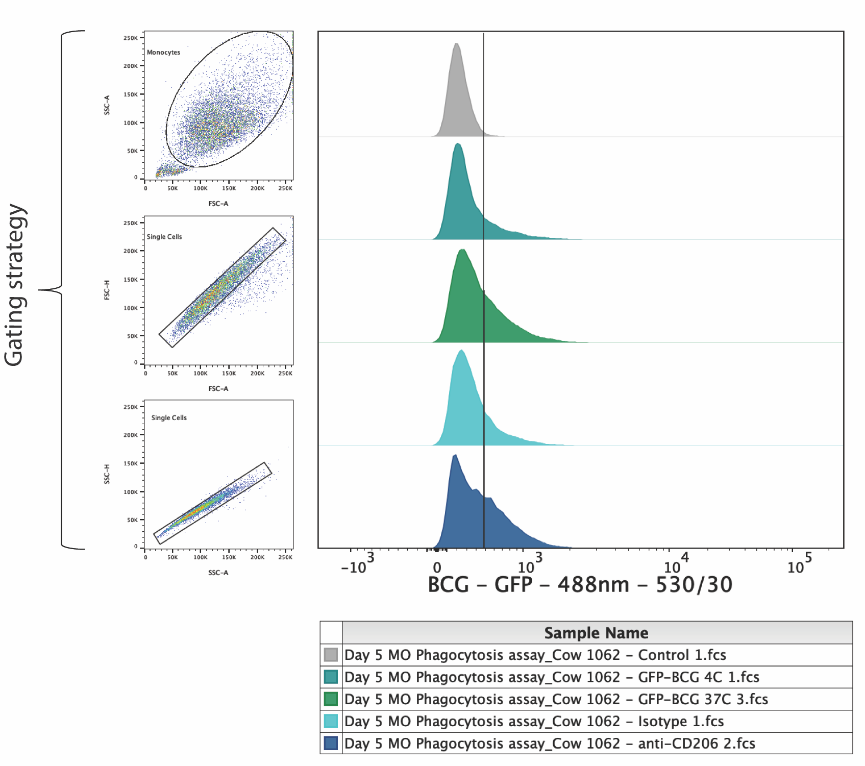

*Supplementary Figure 4 Gating strategy for the analysis of GFP-BCP uptake by bovine MDMØ*

*Bovine Mo were differentiated into MDMØ for five days in culture, incubated with GFP-expressing BCG (MOI of 5) for 90 minutes at either 4°C or 37°C, and analysed by flow cytometry. Detection of all events (FSC vs SSC) was followed by two doublet discriminations (gating strategy) and assessment of GFP-BCG uptake compared to untreated (grey histogram) and antibody-preincubated (blue histograms) cells.*

*Supplementary Table 1: Primers for PCR of bovine MRC1 gene*

| **Primer Name** | **Primer Sequence (5’-3’)** | **Annealing Temp. (°C)** |
| --- | --- | --- |
| Bovine *MRC1*  promotor to 1111bp | FOR: CGCTCTCTTCTCTCCATCCG  REV: CTGCATATGGCCACCACTGA | 65 |
| Bovine *MRC1*  858 to 1941bp | FOR: CAACAGTGGATGGCAGTGGA  REV: TGGACATTTGGGTTCAGGGG | 65 |
| Bovine *MRC1*  1739 to 2924bp | FOR: TCGAGGAAGGCGTTCAGTTC  REV: TCTTTTCGTGCCTCTTGCCA | 65 |
| Bovine *MRC1*  2759 to 3770bp | FOR: TCATCTGCCAGCGCCATAAC  REV: TGGCCCCAGTTCCTTGTAAA | 65 |
| Bovine *MRC1*  3694 to 4368bp | FOR: ACAGAGCACTCAGCATGGAT  REV: TACTAGATGGCCGCATGTTCA | 62 |

*Supplementary Table 2: Antibodies used for staining of bovine Mo and MDMØ*

| **Antibody** | **Clone** | **Manufacturer** |
| --- | --- | --- |
| Mouse anti-bovine CD14: FITC | CC-G33 | Bio-Rad |
| Mouse anti-human CD16: FITC | KD1 | Bio-Rad |
| Mouse anti-human CD163: RPE | EDHu-1 | Bio-Rad |
| Mouse anti-human CD206: PE | 3.29B1.10 | Beckman-Coulter |
| Mouse anti-bovine MHCII DR: RPE | CC108 | Bio-Rad |
| Mouse IgG1 negative control: FITC | Isotype control | Bio-Rad |
| Mouse IgG1 negative control: RPE | Isotype control | Bio-Rad |
